# BaNDyT: Bayesian Network modeling of molecular Dynamics Trajectories

**DOI:** 10.1101/2024.11.06.622318

**Authors:** Elizaveta Mukhaleva, Babgen Manookian, Hanyu Chen, Ning Ma, Wenyuan Wei, Konstancja Urbaniak, Grigoriy Gogoshin, Supriyo Bhattacharya, Nagarajan Vaidehi, Andrei S. Rodin, Sergio Branciamore

## Abstract

Bayesian network modeling (BN modeling, or BNM) is an interpretable machine learning method for constructing probabilistic graphical models from the data. In recent years, it has been extensively applied to diverse types of biomedical datasets. Concurrently, our ability to perform long-timescale molecular dynamics (MD) simulations on proteins and other materials has increased exponentially. However, the analysis of MD simulation trajectories has not been data-driven but rather dependent on the user’s prior knowledge of the systems, thus limiting the scope and utility of the MD simulations. Recently, we pioneered using BNM for analyzing the MD trajectories of protein complexes. The resulting BN models yield novel fully data-driven insights into the functional importance of the amino acid residues that modulate proteins’ function. In this report, we describe the BaNDyT software package that implements the BNM specifically attuned to the MD simulation trajectories data. We believe that BaNDyT is the first software package to include specialized and advanced features for analyzing MD simulation trajectories using a probabilistic graphical network model. We describe here the software’s uses, the methods associated with it, and a comprehensive Python interface to the underlying generalist BNM code. This provides a powerful and versatile mechanism for users to control the workflow. As an application example, we have utilized this methodology and associated software to study how membrane proteins, specifically the G protein-coupled receptors, selectively couple to G proteins. The software can be used for analyzing MD trajectories of any protein as well as polymeric materials.

## Introduction

The dynamics of proteins play a critical role in deciphering their function which in turn is important for designing drugs targeting proteins. Molecular dynamics (MD) simulations are a widely used toolbox to simulate and understand the dynamics of proteins and their complexes ^1–5^. In combination with experimental data from NMR, DEER, and other spectroscopic methods MD simulations provide a detailed atomistic-level view of the protein dynamics and their function ^6–22^. There has been exponential growth in the feasibility of long-timescale MD simulations of proteins and protein complexes that provide rich information on their dynamic properties. At the same time, there is a dire need to shift the usual paradigm of structural bioinformatics from studying single structures to analyzing conformational ensembles.

MD simulation trajectories provide an abundance of high-dimensional data, which makes it a perfect candidate for network-based secondary analysis. Currently, there are multiple network-centric approaches to the analysis of MD simulations data. They can be divided into two main categories: (i) physical proximity based – protein is represented as a graph with residues being nodes, and edges indicating physical interaction between residues with their weight proportional to correlation coefficients extracted from MD simulations ^23–27;^ or (ii) interaction energy based – protein residues make the network’s nodes and edge weights are based on interaction energies between residues, where strong attractive interactions receive higher weights, and weak interactions lead to disconnection ^28–33^. These methods are focused on the residues as variables, and there are currently no network-centric tools that represent residue pairs as network nodes for inferring and quantifying dependencies between inter-residue distances from dynamics data. The existing methodology either simply concentrates on the residues (not residue pairs) or uses dimensionality reduction techniques (primarily principal component analysis, PCA) to analyze the residue pairs data ^34–36^. The latter approach suffers from the inherently low interpretability of PCA, and from the related issue of its inability to separate induced and direct (non-transitive) dependencies. Moreover, there is currently no method based on probabilistic relationships between residues/residue contact pairs that can infer dynamic dependencies in a more data-driven, unbiased manner, regardless of the residues’ spatial proximity. This gap highlights the need for a framework that can go beyond physical proximity and correlations, enabling a deeper understanding of residue-residue relationships within the broader context of protein dynamics. Therefore, we reasoned that Bayesian Network (BN) modeling (BNM) was a natural fit with the MD simulation data, due to the capacity of BNs to infer nonlinear non-transitive dependencies (suggesting directional causality) among different parts of the protein by analyzing dynamical trends. Recently, we developed a workflow to apply BNM to analyze MD simulation trajectories of large protein complexes, namely the G protein-coupled receptors (GPCRs) coupled to trimeric G proteins ^37^. Applying such data-driven interpretable network models led to the identification of the previously unknown important GPCR:G protein interface residue pairs that contribute significantly to G protein coupling strength and to their selectivity/promiscuity ^37^.

Since BNM inherently generalizes to different proteins and other materials where MD simulations are used, in this report we describe the BaNDyT software package that implements the specialized BNM application to MD simulation trajectories, in general. We believe that this software represents the first comprehensive BNM solution for MD simulations, including advanced features not available with other network models. We include a pragmatic description of how to use the software, and how to interpret the results.

## Methodology

### BaNDyT workflow

**Figure 1.**
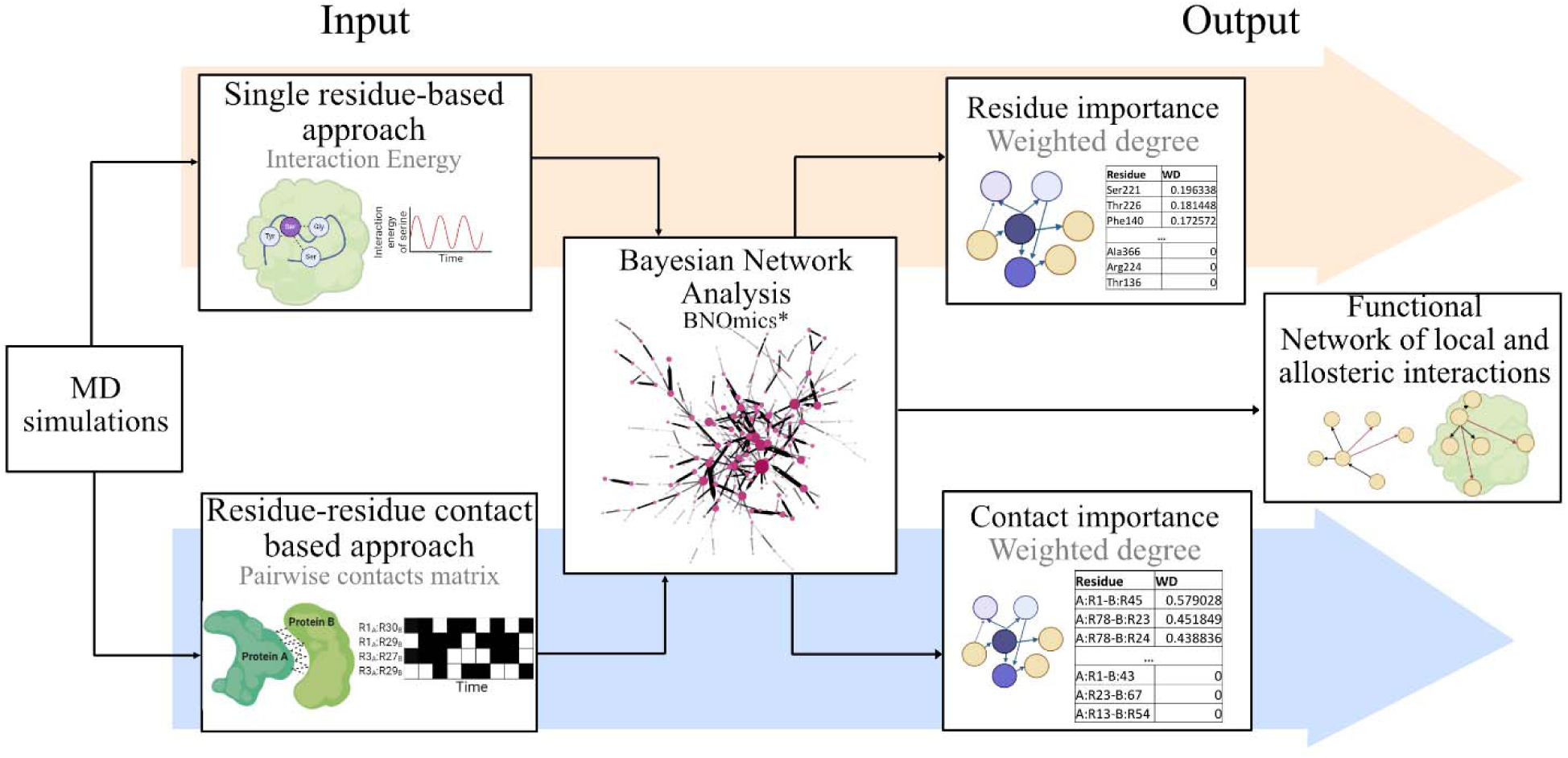
Workflow used in BaNDyT. Step 1 involves performing MD simulations to obtain dynamics trajectories. Step 2 includes extracting data from MD simulation trajectories on single residues and/or pairwise residue-residue interactions as an input for BN analysis. Step 3 uses BNOmics (highly scalable generalist BN modeling software previously developed by us ^38,39^) to construct BN model(s). Step 4 involves visualizing, interpreting, and quantifying the network properties from the BN graph(s) to identify key residues and residue interactions. (BaNDyT can be obtained at GitHub: https://github.com/bandyt-group/bandyt)

#### Step 1: MD Simulations of protein of interest

All-atom molecular dynamics (MD) simulations or any type of enhanced sampling or coarse grain simulations can be performed using standard protocols to generate trajectories that capture the relevant motions and conformational changes ^40–44^. Since the protocols for performing MD simulations vary with the systems simulated, we leave it to the users to formulate their protocols for performing the simulations. To obtain a stable BN downstream, the user should test for the convergence of the MD simulation trajectories. The users could perform multiple MD runs with different starting velocities to ensure robust statistical sampling of the conformational space that would subsequently yield a stable BN. The number of snapshots stored in each trajectory will affect the network model’s robustness – based on our experience, to achieve favorable dimensionality for BNM (given the likely typical protein sizes), we suggest that the user stores snapshots at least every 100ns.

#### Step 2: Generating input variable values for Bayesian Network Modeling

Properties describing each residue in a protein for every MD simulation snapshot can be used as input variables for deriving BN models. Examples of such residue-based properties include but are not limited to (i) interaction of each residue with the rest of the protein, (ii) packing of each residue with its neighboring residues ^45^, (iii) flexibility of each residue, (iv) torsion angles of each residue, or any other such residue-based attributes.

Properties describing the interaction between pairs of residues in every MD snapshot can also be given as input variables to BNM. This is unique to BNM and has not been done before with other network models. Such pairwise residue properties could be the residue contacts or interaction energies between pairs of residues calculated for each MD snapshot. Parameters such as contact frequency, interaction strength, and changes in contacts over time are analyzed to understand the network of interactions within the protein. Pairwise residue properties can be calculated between residues within a protein or in the interface between proteins forming a complex.

Both types of data can be obtained using standard MD analysis tools readily available ^40–44,46–48^. Such input data combined with BN analysis yields a deeper understanding of the dependencies between inter- or intra-protein interactions.

#### Step 3: Constructing Bayesian Networks from Molecular Dynamics data with BaNDyT

The data obtained from the two approaches in Step 2 is then used to construct a BN with the BNOmics software, previously developed by us for highly scalable multi-scale BN modeling ^38,39^. Briefly, given a set of input variables, BN modeling generates a DAG (Directed Acyclic Graph) with these variables as nodes connected by the edges that are direct probabilistic dependencies between the variables. BN modeling is an interpretable unsupervised machine learning network-centered methodology, and the residue-based or contact-based properties input variables correspond to the nodes in the network model. Edge strengths indicate probabilistic dependency strengths that can be estimated via a variety of scoring criteria (BNOmics implements the minimum uncertainty (MU) criterion that is based on the resolution limit as well as the customary AIC and MDL/BIC). The sum of all the edge strengths (evaluated on a universal 0-1 scale when using MU ^38^) of a node provides information on its dependencies on adjacent nodes, in the probabilistic space, irrespective of the nodes’ structural location. The nodes that are connected to a given node in the BN can be structurally local or non-local (allosteric). Such information on allosteric dependencies (i.e., when the nodes are adjacent in the probabilistic space/BN but remote in the physical space) is valuable in determining the allosteric communication mechanism in proteins ^17,18,49–52^. The sum of edge strengths of each node is also known as “node strength,” or “weighted degree”. It should be re-emphasized here that, in contrast with most other network-centered methods, BN modeling aims to filter out spurious dependencies induced by multicollinearities.

### Performing Bayesian Network Modeling of Molecular Dynamic Trajectories data with BaNDyT

Listing 1 shows the script that is used to run BaNDyT. The following steps are necessary to obtain the BN graph:

**Listing 1.**
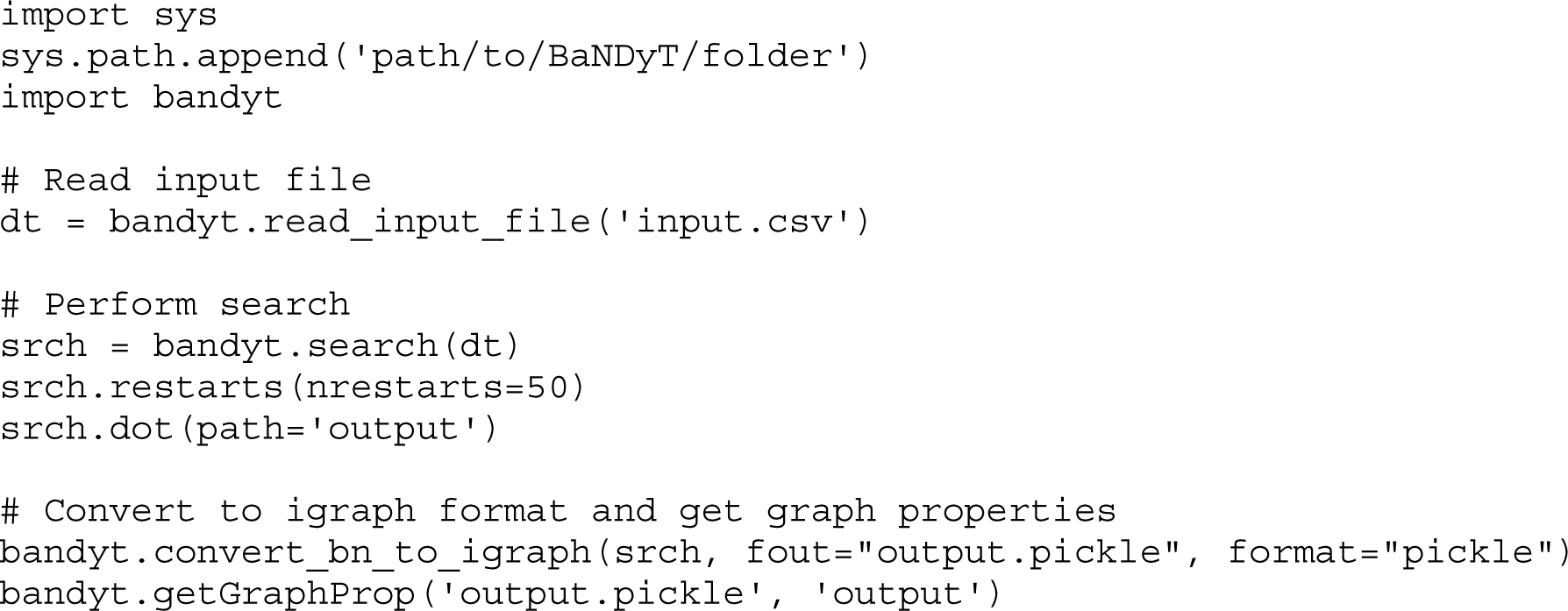
Example of BaNDyT workflow.

#### Data Loading and Discretization

The input data, derived from MD simulation trajectories, is loaded using the bandyt.read_input_file function. In this function, the data containing the continuous variables (for example, residue-centric energy of interaction) is discretized into appropriate bins. By default, BaNDyT utilizes a maximum entropy binning algorithm for discretization ^53^. This approach ensures that each bin contains a roughly equal number of data points, while also maintaining as much randomness as possible in the allocation of data to bins. By maximizing entropy, this method avoids making strong assumptions about the underlying data distribution and ensures that the bins reflect the most uniform possible partition of the data. It is particularly useful in situations where the data distribution is unknown or complex, as it leads to more balanced and informative bins. As the number of observable rare events grows with arity in the case of the recommended scoring function MU, we set the default number of bins as eight, to capture rare events that are still significant.

#### BN search (model selection)

The discretized data is then used to search for the optimal BN topology using the bandyt.search function. There are three scoring functions available in BaNDyT: mdla (based on AIC ^54^), mdlb (based on MDL/BIC ^55^), and mu (based on MU ^38^). We recommend using MU (bandyt.mu) as the scoring function/criterion, due to (i) its superior performance in avoiding false positives (i.e., if the edge, no matter how weak, is in the resulting network, it is highly likely to reflect a real direct dependency) and (ii) edge strengths being estimated on a universal 0-1 scale, which makes downstream comparative analyses possible.

To optimize the number of iterations needed to attain the satisfactory convergence of the BN model selection process, we performed BN reconstruction on proteins of different sizes (Fig. 2A, Table S1). We performed MD simulations on each of these proteins, choosing proteins that vary in sizes from 70 to 1091 residues. We used BaNDyT with MU to generate BN models. We calculated the weighted Hamming distance ^56^ between the graphs to assess the improvement in the network topologies after each restart as compared to the original model to evaluate the similarity. For all the eight proteins of varied sizes considered here, the models appear to show significant improvements up to 40 restarts, after which the changes in topology are minimal (average Hamming distance per node is less than 0.01, Fig. 3B). Therefore, to ensure robustness and correct BN model selection, we recommend repeating the search process with at least 50 restarts (Fig. 2A, B; Fig. S1A).

**Figure 2.**
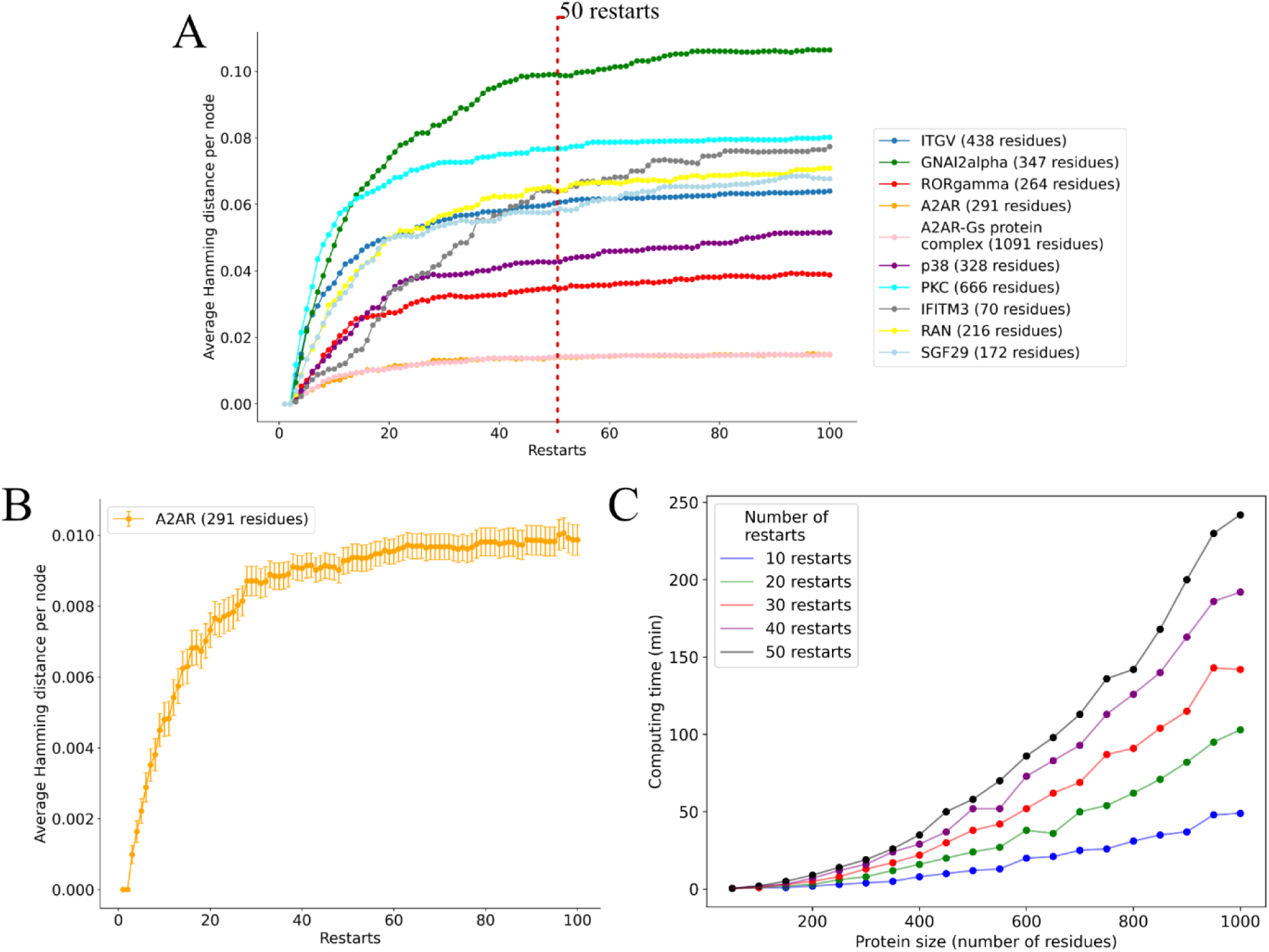
A. Performance and accuracy of Bayesian Network reconstructions for various protein systems. Average Hamming distances per node for BN reconstructions of various proteins with the trajectory length of 5000 frames. Reconstruction for each protein system was replicated 50 times with 100 restarts. The dashed red line indicates the recommended minimal number of restarts. **B.** Average Hamming distances per node for BN reconstructions of A_2A_ receptor. **C.** Computing time of BN models for protein systems of different sizes (on a high-performance cluster with 10G memory and 4 nodes x NVIDIA A100 GPU with 2 x Intel(R) Xeon(R) Platinum 8368 CPU).

**Figure 3.**
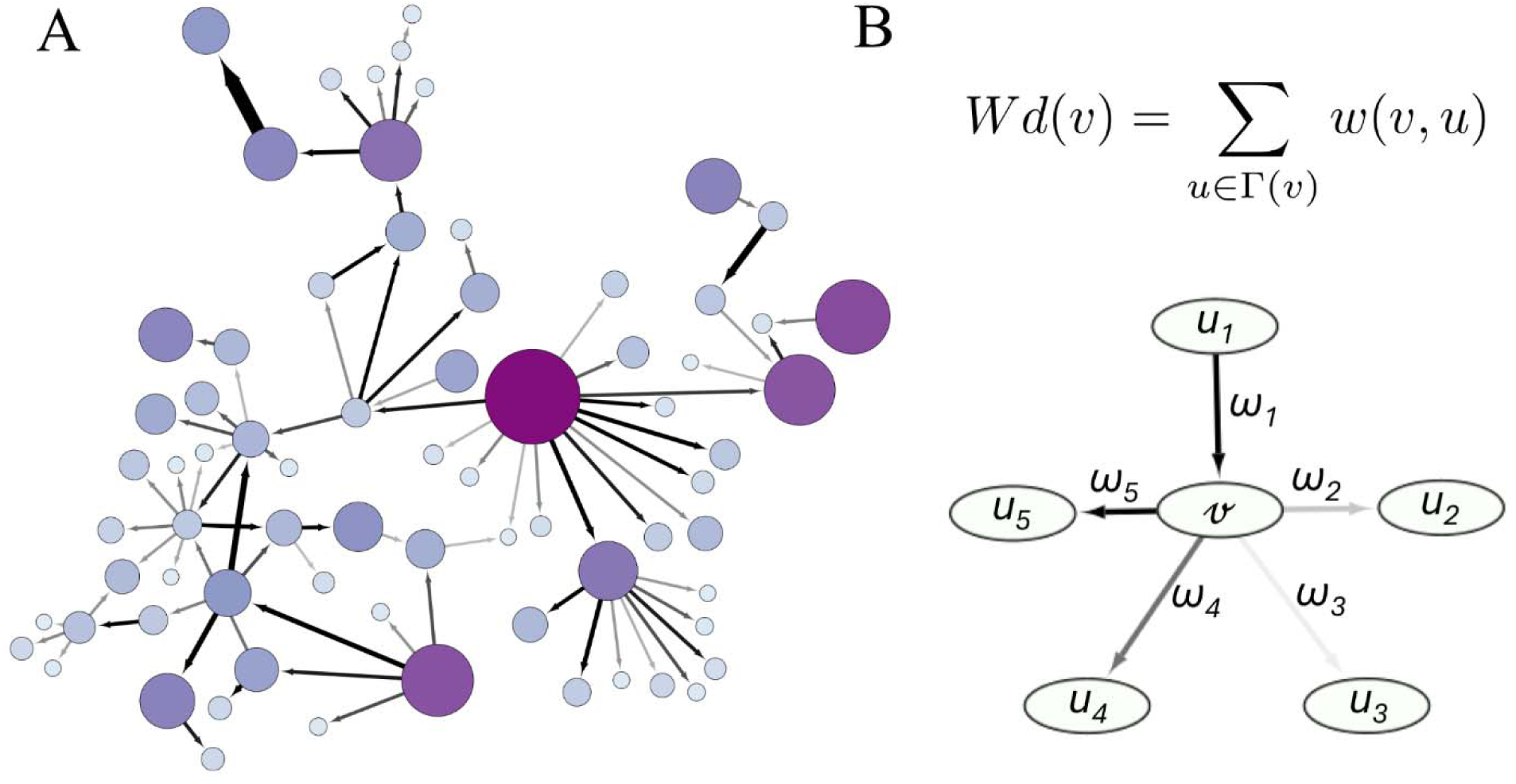
Bayesian Network graph as a main BaNDyT output. **A.** Visual representation of BN graph. The size and color of the node is proportional to the weighted degree of the nodes. The thickness of the arrow is proportionate to the edge weight. **B.** Formula for computing the weighted degree W_d_.

#### Visualizing the BN and Scoring the BN Given the Data

The BN can be visualized using the dot method or using network visualization software such as Cytoscape ^57^, which generates a graphical representation of the network as shown in Fig. 3A. The BN score, obtained with the score_net method, provides a quantitative measure of the network’s fit to the data.

The main BaNDyT output is a BN model which can be represented as a DAG, such as shown in Fig.3A. The graph depicts the probabilistic dependencies between the variables (residues or residue contacts) within the protein. Nodes in the graph represent individual variables, while edges represent the direct (non-transitive, non-spurious) dependencies between them. This representation allows us to infer how changes in one region of the protein might influence other local or distant regions, facilitating the identification of key residues and interactions that are critical for the protein function. As a measure of variable importance, local context network properties associated with nodes (such as betweenness, closeness, etc.), can be extracted. In our case, we have observed that weighted degree shows a strong correlation with experimental data to provide insight into the functional importance of residues and protein-protein interactions ^37^. The weighted degree of node *v* (Fig. 3B) can be computed as

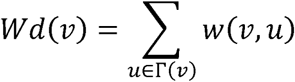

where Γ(*v*) is the set of nodes in the immediate neighbor of the node *v; w(v,u)* denotes the weight of the edge between node *v* and node *u*. The weights reflect the strength or significance of the probabilistic dependencies between the nodes, providing insight into the relative importance of these interactions. In the context of the BNM, the weighted degree of the node helps in identifying how central and “influential” the node is within the local network context. A higher weighted degree indicates that the node has stronger or more significant (or numerous) connections with its neighboring nodes, suggesting that it plays a more central role in maintaining the structure and function of the protein.

#### Step 4: Interpretation of BN Graphs

The final step involves the interpretation of the BN graphs to gain mechanistic insight into the protein dynamics. Here, we aim to translate back from the probabilistic space to the physical space.

- Residue Importance: In the single residue approach, the network can be analyzed to determine the significance of individual residues in the overall structure and function. This is done by evaluating metrics such as weighted degree (Fig. 3B).
- Contact Importance: Similarly, for the pairwise approach, the importance of residue-residue contacts is assessed based on the BN weighted degree. This involves identifying key interactions that are critical for the stability, cooperativity, and function of protein.
- Local and Allosteric Network Interactions: The BN graph provides information (direct node adjacency) on both local interactions (close spatial proximity) and allosteric interactions (distant residues influencing each other). This probabilistic perspective helps in understanding how local changes can propagate through the protein to affect distant sites, shedding light on mechanisms of allosteric regulation.

#### Example of single residue-based analysis

Here, we will use the histamine H_1_ receptor as an example. MD simulation was started from PDB ID 7DFL ^58^, which includes the receptor itself and its coupling partner, trimeric Gq protein. We performed multiple runs of all-atom MD simulations in explicit 1-palmitoyl-2-oleoyl-sn-glycero-3-phosphocholine (POPC) bilayer with cholesterol with a total simulation time of 5µs (see Methods for more details on MD simulation setup). The main input data for the single residue-based approach is the interaction energy for each protein residue (Fig. 4A, Table S2). To obtain this data, we have used *gmx energy* module from GROMACS2022 version ^59^ to calculate the energy of 273 residues of H_1_ receptor. By leveraging this input, a BN was constructed to model the probabilistic dependencies among the residues within the receptor, using previously described parameters (8 bins for discretization of input data, 50 restarts for network convergence; Data S1-2). This network effectively captures intricate interactions and allows for a comprehensive analysis of the residue relationships and their potential impact on receptor function. Upon obtaining the BN graph, we calculated the weighted degree to rank the residues based on their importance within the network (Fig. 4B, C; Table S3). The top 25 percentile of residues, as determined by their weighted degree, are identified as the most important. These key residues were further analyzed to categorize the edges in the network into neighboring interactions and allosteric interactions, based on structural information. Notably, residues within the ligand-binding site were found to have direct connections (in the probabilistic/BN space) with the G protein binding interface and mediator regions, underscoring that minor changes in one location can propagate to distant regions, significantly impacting the receptor’s function (Fig. 4B, C).

**Figure 4.**
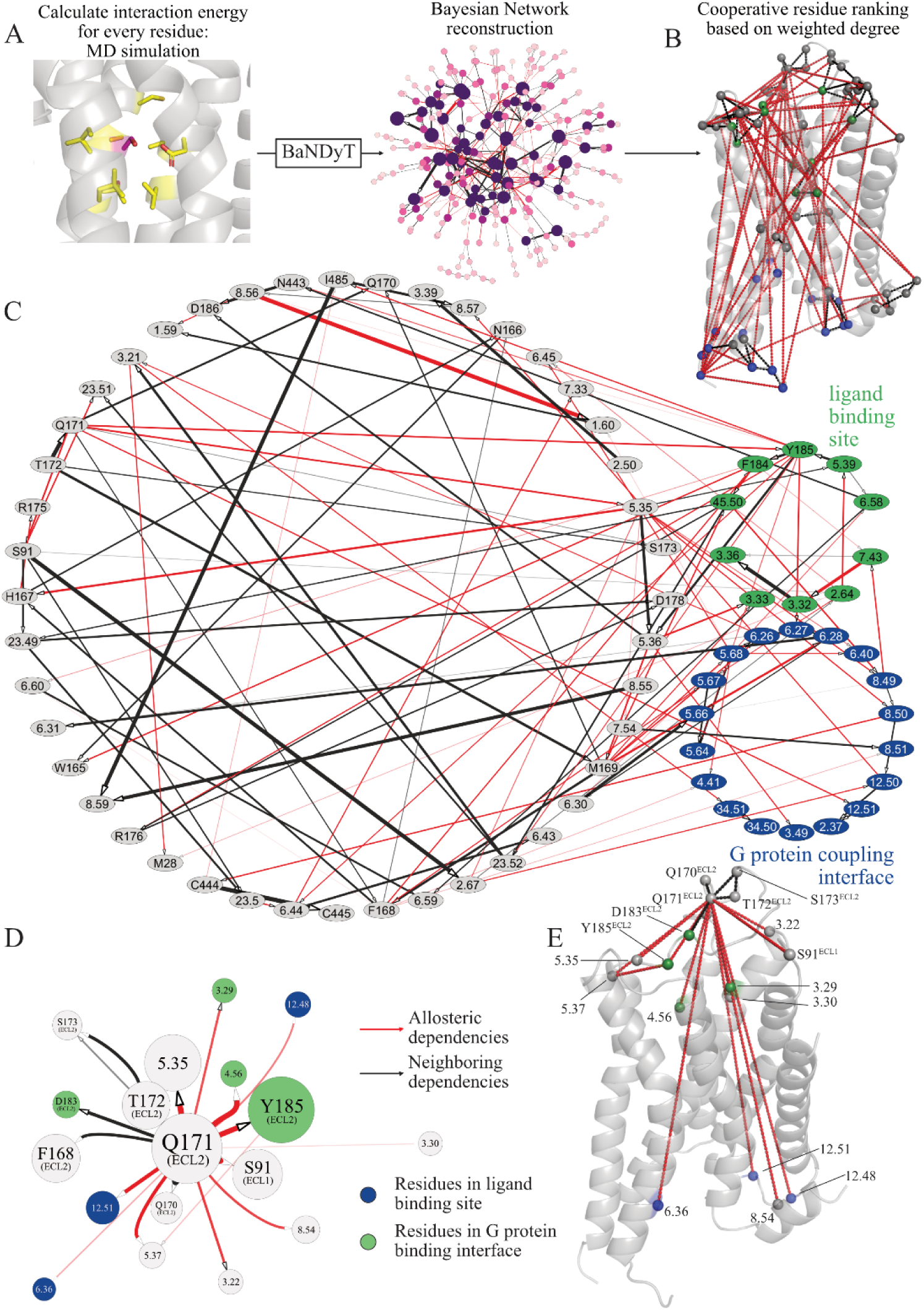
Bayesian Network modeling and structural analysis of H_1_ receptor. **A.** BN modeling of H_1_ receptor network: interaction energies of each residue in H_1_R were used with BaNDyT to obtain the BN graph. **B.** Structural representation of subnetwork of top 25 percentile of nodes based on weighted degree (in C). Ligand-binding site residues are colored green, G protein binding interface residues - blue. Red dash lines represent allosteric probabilistic relationships, black – neighboring relationships. **C.** BN subnetwork of the most important residues (top 25 percentile) based on the weighted degree. Green nodes correspond to ligand-binding site residues, blue nodes – G protein binding interface residues. Red arrows represent allosteric relationships, black – neighboring relationships. **D.** The Markov neighborhood of residue Q171. Size is proportional to weighted degree; arrow thickness is proportional to edge weight. Red arrows represent allosteric relationships, black – neighboring relationships. Green nodes correspond to ligand-binding site residues, blue nodes – G protein binding interface residues. **E.** Structural representation of the network depicted in D.

One of the most important aspects of BN graph analysis is the ability to analyze a community of residues that have established probabilistic relationships. Fig. 4D, E presents an example of the Markov neighborhood (an immediate-adjacency subset/simplification of the Markov blanket) of an important node, residue Q171 in extracellular loop 2. Markov neighborhood of a node refers to the set of all its adjacent nodes (neighbors); the state of the given node is conditionally independent of all other nodes in the graph, given the states of the nodes in its Markov blanket. As seen from the network and structural representation (Fig. 4C, D), the network relationships of this residue can be separated into neighboring and allosteric. Although this residue is not part of the immediate ligand pocket, it evidently influences other functionally important residues, such as 6.36 (G protein binding residue) and 4.56 (ligand binding site residue, mutations that affect protein function ^60^). Thus, the BN model provides valuable insights into how specific residues, even the ones that are not located in the functionally important region, can play pivotal roles in the receptor’s functionality through their influence on the network of interactions.

#### Example of pairwise contact-based analysis

Here, we will use the muscarinic receptor 1 (M_1_) coupled to the Gq protein as an example. Similarly to H_1_ receptor, we have performed all-atom MD simulations of GPCR:G protein complex (PDB ID: 6OIJ ^61^) in POPC membrane according to the protocol, as described in Methods. To investigate the protein-protein residue contacts within this complex, we utilized GetContacts ^62^, which provided a detailed map of these interactions (Fig. 5A). GetContacts-generated output was converted into a binary matrix of residue contacts fingerprints, where 1 represents presence of the contacts in the trajectory frame, and 0 represents absence. Unlike conventional methods, we did not apply any frequency cutoffs, as BN analysis is capable of capturing significant rare events (infrequent contacts) from transient interactions (recall that the MU criterion is optimized for guarding against false positives). This resulted in the dataset of 427 residue-residue contacts with 25,000 trajectory frames as an input for BaNDyT (Table S4). BaNDyT was run using default parameters of 50 restarts, while discretization was not needed due to data already being binary. The primary outcome of our BN analysis was the BN graph, illustrating the direct dependencies between variables—in this case, the residue contacts (Fig. 5B; Data S3-4). The weighted degree of each node was used as a measure of importance within the network, reflecting the cooperativity of protein-protein interactions observed throughout the MD simulation trajectory (Table S5).

**Figure 5.**
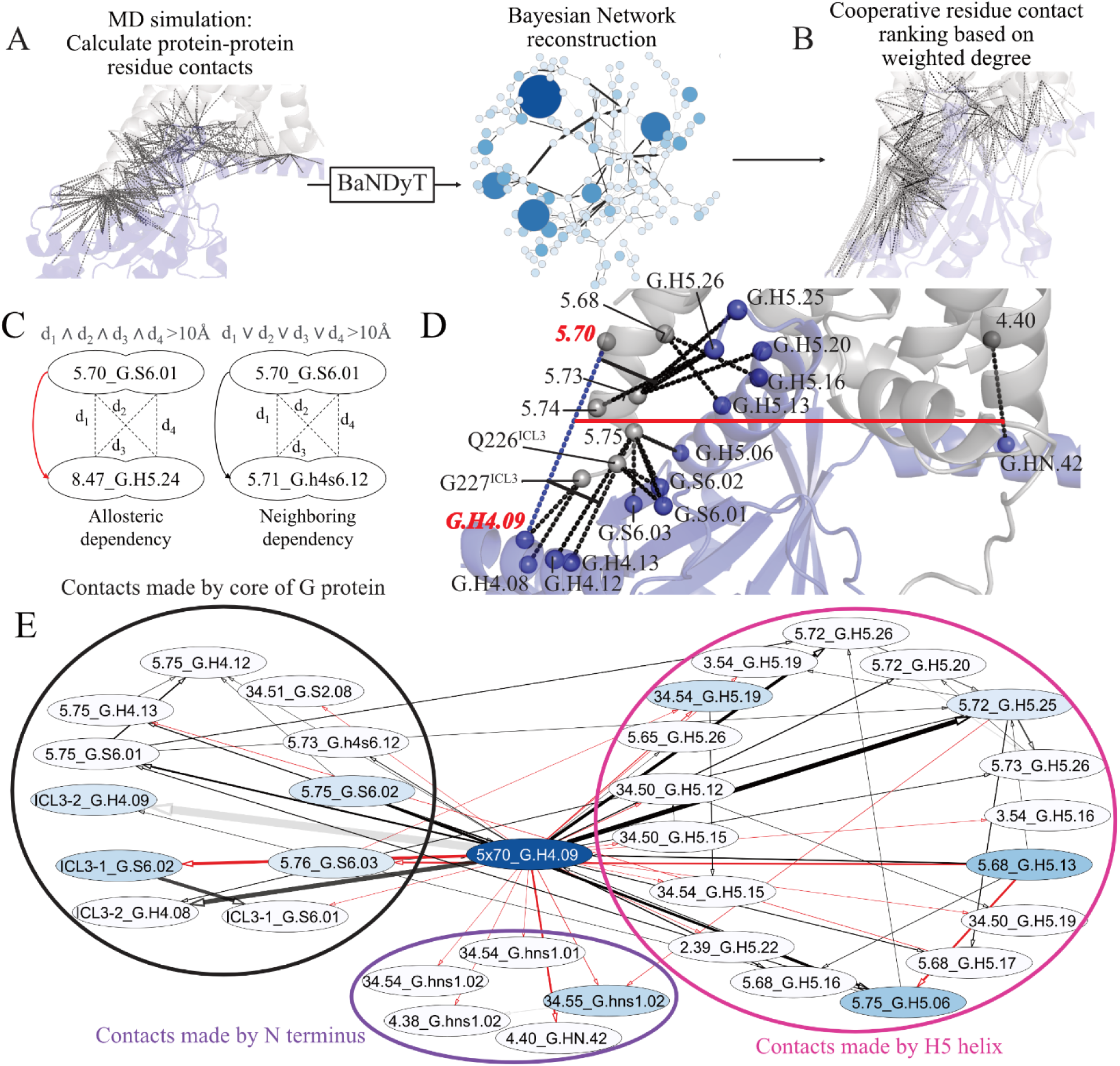
Bayesian Network modeling and structural analysis of M_1_:Gq protein residue contact network. **A.** BN modeling of M_1_:Gq protein contact network: contact fingerprints were used with BaNDyT to obtain the BN graph. **B.** BN graph of M_1_:Gq protein residue contact network. Size and color intensity are proportional to the weighted degree of the node, thickness of the arrows is proportional to the edge strength. **C.** Schematic representation of neighboring and allosteric dependencies. Nodes (GPCR:G protein residue contacts) are represented by oval shapes. The pairwise distance between nodes is marked as a black dashed line. A node dependency is considered as neighboring if at least one of the distances (d_1_, d_2_, d_3_, d_4_) is less than 10Å and allosteric if all distances (d_1_ to d_4_) are larger than 10Å. **D.** Structural representation of nodes (contacts) that are connected to node 5.70_G.H4.20 in the BN graph with edge weight higher than 0.01. M_1_ is represented as grey cartoon, Gq protein – as blue cartoon. Cα atoms of residues involved in the contacts are shown in spheres, black dash lines represent contacts. Red solid line shows allosteric dependency, black solid line – neighboring. **E.** The Markov neighborhood of contact 5.70_G.H4.09. The nodes are grouped by their location in the G protein structure: N terminus, core, or H5 helix. Node color is proportional to weighted degree; arrow thickness is proportional to edge strength. Red arrows represent allosteric relationships, black – neighboring relationships.

We identified the top 25 percentile of nodes with the highest weighted degrees as the most cooperative interactions, which were subsequently analyzed in the context of protein biology (Fig. 5B, Table S5). This approach has been previously applied to the analysis of six different GPCR protein complexes, revealing significant differences in how various G protein coupling types contribute to the cooperativity of these interactions and highlighting the prevalence of certain types of contacts across different coupling types ^37^.

Recall that the BN graph connects variables in the probabilistic space without considering their structural positions, meaning that contacts that are not in close physical proximity can still be directly connected within the network. This means that even distant contacts may influence each other during the MD trajectory. To further investigate these interactions, we categorized BN dependencies into two groups: neighboring dependencies, where the Cα atoms of the GPCR and Gα subunit residues in two contacts are within 10Å of each other, and allosteric dependencies, where these atoms are more than 10Å apart (Fig. 5C).

Both allosteric and neighboring dependencies contribute to the importance of a node (contact) within the network. For example, in Fig. 5D, E, we explore the Markov neighborhood of the most cooperative contact in the M_1_ protein complex, 5.70_G.H4.20. While this contact is positioned in the core region of the G protein, it has multiple interactions with the contacts formed by H5 helix in the network graph (Fig. 5E). Given the established importance of H5 helix interactions in G protein coupling, disruptions to contact 5.70_G.H4.20 may have profound effects on the dynamics of dependent H5-helix contacts, potentially compromising GPCR-G protein interaction. Our prior research highlights the core’s heightened influence in Gs proteins over Gq proteins ^37^. As the M_1_ complex comprises a Gq protein chimera with a Gs protein core ^61^, this configuration could manifest Gs-like characteristics in the interplay between the core and the H5 tip.

### Concluding remarks

Here, we have presented BaNDyT, a novel Bayesian Network Modeling software specifically designed for the analysis of MD simulation trajectories. By leveraging the power of BNM, BaNDyT provides an intrinsically interpretable, scalable and fully data-driven unsupervised machine learning approach to uncovering functional relationships between residues and residue pairs in proteins, going beyond traditional MD analysis methods. Our approach addresses several limitations in the current network-based methodologies, such as their reliance on residue-centric variables and the surfeit of spurious dependencies. With the ability to capture both local and allosteric dependencies, BaNDyT provides a comprehensive framework for analyzing the dynamic interplay within proteins and their complexes, as demonstrated through its application to GPCRs and their selective coupling to G proteins. Furthermore, BaNDyT’s versatile Python interface makes it a robust tool for the broader scientific community to explore MD simulation data across various systems, including polymeric materials and other complex biomolecular structures. The resulting Bayesian networks offer new insights into key residues and interactions that are critical for protein stability, function, and allosteric regulation, paving the way for more targeted and effective therapeutic interventions.

## Methods

### Preparation of Protein Structures for MD simulations

All protein and protein complexes were prepared for MD simulations using the corresponding experimental structures: A_2A_R:Gs protein complex – 6GDG^63^, H_1_R-Gq protein complex – 7DFL^58^, M_1_R-Gq protein complex - 6OIJ^61^, SGF29 – 3ME9^64^, ITGV – 3IJE^65^, RORgamma – 4WLB^66^, PKC – 3IW4^67^, GNAI2 - 6CRK^68^, RAN - 2MMC^69^. Mutations in structures were reverted to their wild-type residues using Maestro (Schrödinger Release 2020-1: Maestro, Schrödinger, LLC, New York, 2020). Ligands were parameterized using ParaChem (https://cgenff.umaryland.edu). Missing side chains and loops (fewer than 5 residues absent) were modeled into the 3D structures proteins. Residues within 5 Å of mutation sites were minimized using MacroModel, with position restraints applied to all backbone atoms. Protein termini were capped with neutral acetyl and methylamide groups, and histidine protonation states were assigned via the Maestro protein preparation wizard. In case of A_2A_R and H_1_, receptors were embedded into an explicit POPC bilayer membrane using the PPM 2.0 function from the Orientation of Proteins in Membranes (OPM) tool ^70^, and hydrated with TIP3P water containing 0.15 M NaCl using CHARMM-GUI ^71,72^. The final system dimensions were approximately 120 Å × 120 Å × 170 Å, and all systems were characterized using the CHARMM36m force field ^73^.

### MD Simulations for Proteins and Protein complexes

All molecular dynamics simulations were conducted using the GROMACS 2019 ^48^ package with a 2 fs integration timestep. The prepared systems underwent energy minimization with position restraints of 10 kcal/mol·Å² applied to the heavy atoms of the proteins, ligand, and lipids if present. A 1-ns heating phase followed, gradually raising the temperature from 0 K to 310 K under the NVT ensemble with the Nosé-Hoover thermostat. This was followed by equilibration in the NPT ensemble, where the initial 1-ns run retained the 10 kcal/mol·Å² restraints, which were then reduced incrementally (from 10 to 5, and finally to 1 kcal/mol·Å²) in 5-ns steps. The final equilibration phase was a 50-ns simulation without position restraints. The last snapshot of this equilibration served as the starting point for five production runs, each 1 μs in length, initiated with randomly generated velocities. The pressure was controlled at 1 bar using the Parrinello-Rahman method ^74^. Nonbonded interactions were calculated with a 12 Å cutoff, and long-range interactions were handled using the Particle Mesh Ewald (PME) method ^75^. The LINCS algorithm constrained all bonds and angles of water molecules.

### Calculation of Residue Interaction Energy

Ensemble trajectories were used to analyze interaction energies between protein residues. Interaction energies for each residue with the rest of the protein were calculated using the GROMACS “ energy” tool ^48,59^. The total nonbonded interaction energy, comprising short-range (within 12 Å) coulombic and van der Waals forces, was extracted from the energy log file and summed to obtain the overall nonbonded interaction energy.

### Calculating Fingerprints of Pairwise Interactions Between M_1_R and Gαq Protein

Pairwise residue contacts between M_1_R and Gq protein were analyzed using the Python script library “GetContacts” ^62^ (https://www.github.com/getcontacts). This tool was employed to identify different types of interactions, including salt bridges (cutoff < 4.0 Å), hydrogen bonds (cutoff < 3.5 Å with an angle < 70°), van der Waals contacts (difference < 2 Å), π-stacking interactions (distance < 7.0 Å with an angle < 30° between aromatic planes), and cation-π interactions (distance < 6.0 Å with an angle < 60°). The analysis was conducted over the 5μs trajectory, excluding lipids, water and ions. The atom selection groups were aligned with the relevant residues from both the GPCR and the G protein α-subunit. Each residue was mapped to its corresponding generic residue number using BW numbering for the GPCR and Common G protein numbering for G proteins. Transmembrane helix ends were adjusted using BW numbering. Custom Python scripts were used for one-hot encoding, generating binary contact fingerprints for each simulation frame, where “1” indicates a contact and “0” represents its absence.

## Supplementary Material

A Python-based Jupyter Notebook that contains a tutorial for BaNDyT is provided in the supplementary material and can be accessed at https://github.com/bandyt-group/bandyt-tutorial.

## Data and Software Availability

BaNDyT is available on GitHub at: https://github.com/bandyt-group/bandyt. A demo with instructions on how to use the software is available on GitHub at: https://github.com/bandyt-group/bandyt-tutorial. An XLSX file containing interaction energy data and a CSV file with a residue-residue contact matrix, both derived from molecular dynamics simulations and used as inputs for the BaNDyT analysis, are provided in the Supplementary Information (SI). Outputs of BaNDyT for showcases are provided as CSV (network properties), GRAPHML and PICKLE files (graphs) in the SI. Full simulation data will be provided by the authors upon request.

## Funding / Acknowledgement

This work was funded by grants from the National Institutes of Health R01-GM117923 to N. V., R01-LM013876 to N. V., A. R., Sergio Branciamore, and Supriyo Bhattacharya, and R01-LM013138 to A. R. The content is solely the responsibility of the authors and does not necessarily represent the official views of the National Institutes of Health. Additional support is acknowledged by Dr Susumu Ohno Chair in Theoretical Biology (held by A. R.), and Susumu Ohno Distinguished Investigator Fellowship (to G. G.).

