## Supplemental Information for "BaNDyT: Bayesian Network modeling of molecular Dynamics Trajectories"

**This PDF file includes:**

Figure S1

**Other Supporting Information for this manuscript include the following:**

Jupyter Notebook Tutorial for BaNDyT

Table S1-5

Data S1-4

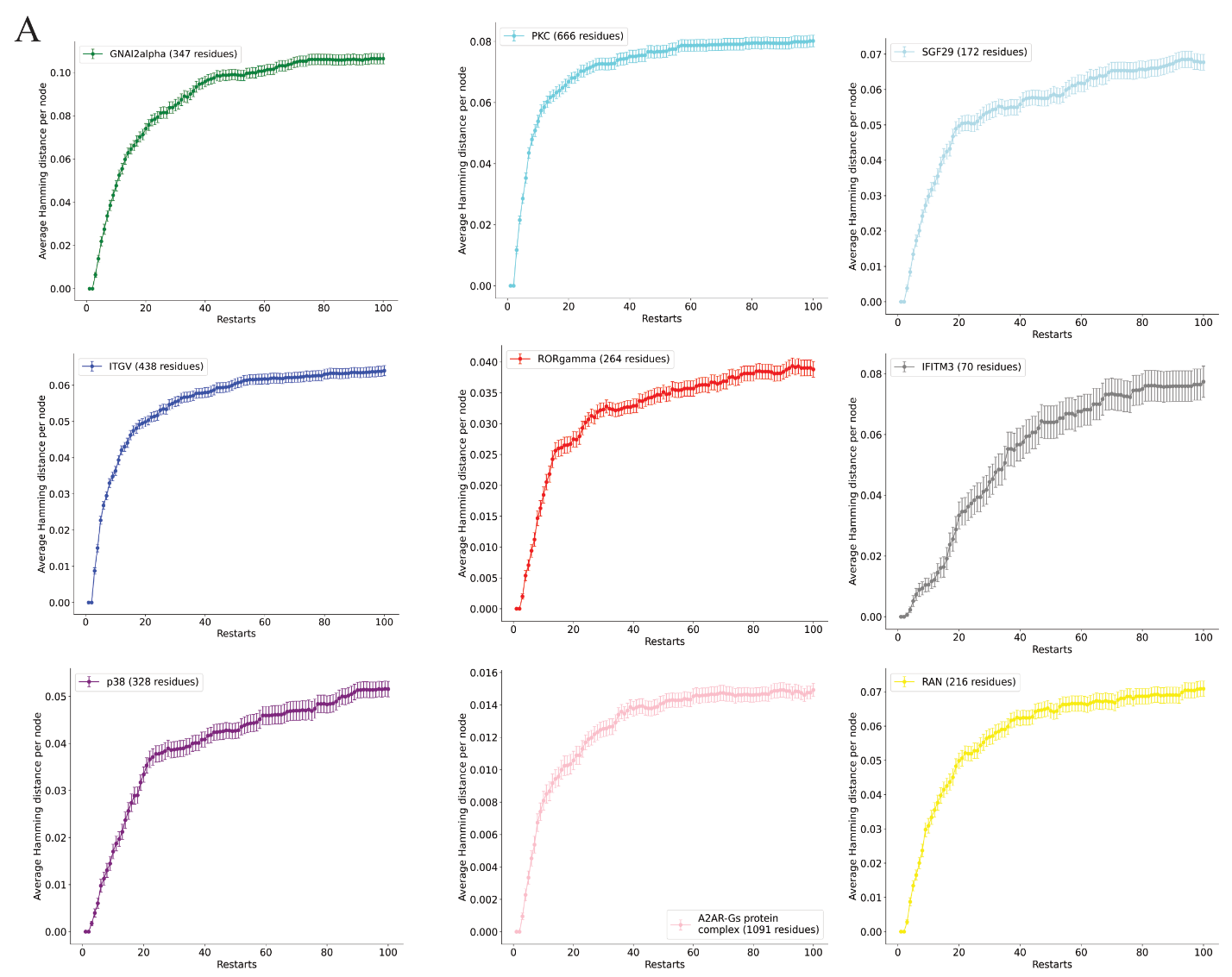

**Supplementary Figure S1. Convergence of Bayesian Network reconstructions for various protein systems. A.** Average Hamming distance per node for following proteins: top row (left to right) – Gαi2 protein, protein kinase C (PKC), SAGA complex associated factor 29 (SGF29); middle row – integrin V (ITGV), RAR-related orphan receptor gamma (RORγ), interferon induced transmembrane protein 3 (IFITM3); bottom row - p38 mitogen-activated protein kinase (p38), A2AR-Gs protein complex, ras-related nuclear protein (RAN).
